# Absence of muricholic acid due to Cyp2c-deficiency protects against high fat diet-induced obesity in male mice but promotes liver damage

**DOI:** 10.1101/2021.05.10.443487

**Authors:** Antwi-Boasiako Oteng, Sei Higuchi, Alexander S. Banks, Rebecca A. Haeusler

**Affiliations:** Naomi Berrie Diabetes Center, Columbia University Medical Center, New York, NY, USA; Department of Pathology and Cell Biology, Columbia University Medical Center, New York, NY, USA; Division of Endocrinology, Beth Israel Deaconess Medical Center and Harvard Medical School, Boston, MA, USA

## Abstract

**Objective:** Murine-specific muricholic acids (MCAs) are reported to protect against obesity and associated metabolic disorders. However, the response of mice with genetic depletion of MCA to an obesogenic diet has not been evaluated. We used Cyp2c-deficient (Cyp2c^−/−^) mice, which lack MCAs and thus have a human-like bile acid (BA) profile, to directly investigate the potential role of MCAs in diet-induced obesity.

**Methods:** Male and female Cyp2c^−/−^ mice and wild-type controls were fed a standard chow diet or a high fat diet (HFD) for 18 weeks. We measured BA composition from a pool of liver, gallbladder, and intestine, as well as weekly body weight, food intake, lean and fat mass, systemic glucose homeostasis, energy expenditure, intestinal lipid absorption, fecal lipid, and energy content.

**Results:** Cyp2c deficiency depleted MCAs and caused other changes in BA composition, namely a decrease in the ratio of 12α-hydroxylated (12α-OH) BAs to non-12α-OH BAs, without altering the total BA levels. While wild-type male mice became obese after HFD-feeding, Cyp2c^−/−^ male mice were protected from obesity and associated metabolic dysfunctions. Cyp2c^−/−^ male mice also showed reduced intestinal lipid absorption and increased lipid excretion, which was reversed by oral gavage with the 12α-OH BA, taurocholic acid. Cyp2c^−/−^ mice also showed increased liver damage, which appeared stronger in females.

**Conclusion:** MCA does not protect against diet-induced obesity but may protect against liver injury. Reduced lipid absorption in Cyp2c-deficient male mice is potentially due to a reduced ratio of 12α-OH/non-12α-OH BAs.

## 1. INTRODUCTION

Bile acids (BAs) are catabolites of cholesterol that are synthesized in the liver and secreted into the gut. Liver-derived primary BAs are exported as glycine or taurine conjugates, the former most prominent in humans and the latter most prominent in mice, and can be converted to unconjugated and secondary BAs in the gut by resident microbes [1,2]. In the gut, BAs facilitate the absorption of fats, cholesterol, and fat-soluble vitamins. BAs also possess endocrine functions by serving as ligands for the transcription factor farnesoid X-receptor (FXR) and Takeda G-protein coupled receptor (TGR5), thus enabling BAs to play important roles in physiological regulation of lipid and glucose metabolism [3–5].

Human studies have shown that BA levels, composition, and signaling are associated with cardiometabolic diseases such as obesity, dyslipidemia and diabetes [6], highlighting the potential of BA modulation in treating or preventing such disorders. Obesity in humans is positively associated with increased synthesis of BA, impaired serum BA fluctuations and increased synthesis of 12α-hydroxylated BAs, which include cholic acid (CA) and its secondary BA derivative deoxycholic acid (DCA) [7]. Additionally, the ratio of 12α-hydroxylated to non-12α-hydroxylated BAs is positively correlated with human insulin resistance [8]. To establish whether alterations of BA metabolism play a causal role in cardiometabolic disorders, mechanistic studies in pre-clinical models are important. Many such mechanistic investigations have been conducted in rats and mice. However, the use of these models is translationally challenging, because mice produce muricholic acids (MCAs) that are not present in healthy adult humans. MCAs are trihydroxylated at carbons 3, 6, and 7, and their unique structures are thought to cause differential functionality. For one thing, some MCAs can antagonize, rather than activate, FXR [9–11]. Furthermore, MCAs are thought to be poor at promoting fat and cholesterol absorption. This concept is supported by studies that used dietary interventions to increase MCAs in mice, which results in reduced cholesterol absorption [12]. Other studies have indirectly increased MCAs by deleting the 12α-hydroxylase Cyp8b1, thus eliminating the 12α-hydroxylated BAs such as CA and DCA, and increasing the non-12α-hydroxylated BAs (which, in mice, are primarily MCAs). This results in reduced absorption of dietary fat and cholesterol, lower body weight, and protection against diet-induced obesity [13–17]. Thus, it has been suggested that MCAs play a causal role in protection against weight gain and diet-induced obesity. However, the direct effects of MCAs on obesity and associated metabolic disorders has not been evaluated in a mouse model with specific genetic elimination of MCAs.

An insightful study by Takahashi et al [18] demonstrated that the presence of MCAs in mice requires the *Cyp2c* cluster of genes, which includes 14 gene isoforms. Recent studies show that specifically deleting the *Cyp2c70* isoform results in partial or complete reduction of MCAs, with concomitant increase in the levels of chenodeoxycholic acid (CDCA), the major precursor BA for MCA synthesis [19–21]. To determine the effects of eliminating MCAs on fat absorption and susceptibility to obesity and its associated metabolic complications, we used mice with a germline deletion of the 14 Cyp2c isoforms that are clustered on mouse chromosome 19 (Cyp2c^−/–^) [22,23]. BA analysis confirmed the lack of MCAs, and one might predict this to enhance intestinal lipid absorption and increase susceptibility to diet-induced obesity. On the other hand, Cyp2c^−/–^ mice also showed a significant reduction in levels of 12α-OH BAs compared to control mice. Because 12α-OH BAs promote fat absorption [14,16], this suggests the opposing possibility that Cyp2c^−/–^ mice would be less susceptible to diet-induced obesity. To distinguish between these possibilities, we set out to investigate the response to an obesogenic diet by Cyp2c^−/–^ mice.

## 2. MATERIALS & METHODS

### 2.1. Animal model and Interventions

#### 2.1.1. Generation and source of Cyp2c^−/−^ mice

Animal experiments were conducted in wild-type (WT) and Cyp2c-knockout (Cyp2c^−/−^) mice with C57BL/6N background obtained from Taconic Biosciences (New York, NY, USA). The mice (model # 9177) were originally generated by a Cre-mediated germline deletion of 14 out of 15 isoforms of the Cyp2c genes. The deleted genes, which are located within a 1.2 Mb cluster on chromosome 19, include *Cyp2c55, 2c65, 2c66, 2c29, 2c38, 2c39, 2c67, 2c68, 2c40, 2c69, 2c37, 2c54, 2c50, 2c70. Cyp2c44*, which is physically located outside of this cluster, has not been deleted. The resultant chimeras were backcrossed to C57BL/6N WT mice [22]. Male and female heterozygotes were subsequently purchased from Taconic Biosciences and bred to generate knockout mice and littermate controls for this study. Mice were genotyped using a genotyping kit (#KK5621, KAPA Biosystems). The DNA products after PCR were resolved by electrophoresis on a 2% agarose gel with 0.05% ethidium bromide. The primer sequences used for genotyping are provided in Supplementary Table 1.

#### 2.1.2. Chow and high fat diet (HFD) intervention

Male WT (n=7), male Cyp2c^−/−^ (n=7), female WT (n=7), and female Cyp2c^−/−^ (n=7) mice were weaned onto and maintained on standard chow diet (3.4 kcal/g, Purina 5053, 24.7% kcal from protein, 62.1% carbohydrate and 13.2% fat) *ad libitum*. Body weights were measured once every week until mice were euthanized at ~30 weeks old. When mice were between 17-20 weeks old, oral glucose tolerance tests (OGTT) and insulin tolerance tests (ITT) were performed. During the HFD intervention, male WT (n=7), male Cyp2c^−/−^ (n=7), female WT (n=5), and female Cyp2c^−/−^ (n=5) mice were weaned onto standard chow diet until mice were transferred onto a HFD (60% kcal fat, D12492, Research Diets Inc., New Brunswick, NJ, USA) at 7-8 weeks old. The mice were fed the HFD for 18 weeks. Body weights were measured once every week. OGTT and ITT were performed after 12 and 14 weeks on the HFD, respectively.

In both chow and HFD groups, mice were singly housed two-weeks prior to euthanasia, to collect feces and to measure food intake using feeding dispensers that were placed inside the cage. One week before euthanasia, mice were subjected to a body composition assessment for lean and fat mass using time domain NMR (Minispec Analyst AD; Bruker Optics) [24] followed in male mice by indirect calorimetry in metabolic cages for energy expenditure measurement.

Mice were euthanized using CO_2_ asphyxiation followed by cervical dislocation. Blood was collected through the inferior vena cava into EDTA-coated tubes to obtain plasma followed by excision of liver, gonadal white adipose tissue (gWAT), brown adipose tissue (BAT), and intestine for analysis.

All experiments were approved by, and conducted in accordance with the guidelines of the Columbia University Institutional Animal Care and Use Committee.

#### 2.1.3. Glucose tolerance test (GTT)

Following a 6 hour fast, baseline glucose measurements representing timepoint 0 were recorded via tail vein bleeding using OneTouch glucose monitor and strips (LifeScan). Mice were then orally gavaged (in case of OGTT) or injected intraperitoneally (in case of IPGTT) with glucose (2 g/kg body weight) followed by glucose measurements at timepoints 15, 30, 60, 90 and 120 mins.

#### 2.1.4. Insulin tolerance test (ITT)

Following a 5 hour fast, baseline glucose measurements representing timepoint 0 were recorded via tail vein bleeding using OneTouch glucose monitor and strips (LifeScan). Mice were then injected intraperitoneally with insulin (0.5U/kg body weight) followed by glucose measurements at timepoints 15, 30, 45 and 60 mins.

#### 2.1.5. Metabolic cages

Indirect calorimetry and activity measurements were performed with Comprehensive Laboratory Animal Monitoring system (CLAMS) open-circuit Oxymax system (Columbus Instruments). Mice were individually housed for 7 days in metabolic cages. The animals were housed at 22°C for the duration of the measurements with a 07:00-19:00 light photoperiod. Mice were aged 26-29 weeks and 24-25 weeks for experiments on chow or HFD, respectively. Analysis of energy expenditure was performed using CalR [25]. Mice found to have not eaten for more than 36 hours due to a malfunction in the food hopper were excluded from the analysis.

#### 2.1.6. Fasting Re-feeding experiment

Mice maintained on chow diet were fasted overnight, and re-fed for 2 hours with HFD. Blood was collected after fasting and after re-feeding and centrifuged at 2,500 g for 15 mins to obtain plasma.

#### 2.1.7. Taurocholic acid (TCA) gavage experiment

Mice were orally gavaged with 17 mg/kg TCA in 1.5% NaHCO_3_ at 6 pm each day, for 3 consecutive days. The 17 mg/kg dose was chosen based on earlier publications demonstrating that a dose in this range has does not alter the total BA levels but raises the TCA levels in mice [26,27]. On day 4, mice underwent the radiolabeled triolein experiment to measure lipid absorption.

#### 2.1.8. Radiolabeled triolein experiment

For analysis of lipid absorption, mice maintained on chow diet underwent a 4 hour fast between 7 am and 11 am, injected intraperitoneally with poloxamer 407 (1 g/kg body weight; BASA) in PBS [28], then orally gavaged with olive oil (10 μL/g body weight) containing 2.5 μCi of [^3^H]triolein ([9–10-3 H(N)]triolein; PerkinElmer), which is a radiolabeled triglyceride. Mice were bled before the gavage at time 0, and then at 1, 2, 4, 8, and 24-hours post gavage. Blood samples were analyzed for radioactivity (Tricarb 2910TR Scintillation Counter; PerkinElmer).

### 2.2. Bile acid (BA) profile

Liver, gallbladder, and small intestine were doubly homogenized in 50% methanol using a rotor-stator homogenizer, followed by a Dounce Teflon-glass homogenizer. Deuterated BA standard (20 μL of 25 μM d4-cholic acid) were added to 200 μL of each sample and calibrator curves were generated of each BA in charcoal-stripped tissue. To each sample/calibrator, 2 mL of ice-cold acetonitrile was added, then samples were vortexed for 1 hour at 2,000 rpm and centrifuged for 10 min at 11,000 g. Supernatants were transferred to clean glass tube and dried down at 45°C under nitrogen. Each sample/calibrator was extracted a second time in 1 mL of ice-cold acetonitrile, vortexed for 1 hour at 2,000 rpm, and centrifuged for 10 mins at 11,000 g. The supernatant of the second extraction was combined with the first and dried down at 45°C under nitrogen. Each sample was resuspended in 200 μL of 55:45 (vol/vol) methanol:water, both with 5 mM ammonium formate. Samples were centrifuged in UltraFree MC 0.2-μm centrifugal filters (Millipore) and transferred to liquid chromatography-mass spectrometry (LC-MS) vials, and 10 μL were injected into ultraperformance liquid chromatography-tandem mass spectrometry vials (UPLC-MS/MS; Waters).

### 2.3. Plasma analysis

Blood glucose was measured in mice that were fasted for 5 hours by tail vein bleeding using OneTouch glucose monitor and strips (LifeScan). Plasma was obtained after centrifuging blood collected in EDTA-coated tubes for 15 mins at 2,500 g. Plasma insulin was measured using an ELISA kit (# 10-1247-01, Mercodia) according to manufacturer’s protocol. Plasma lipids were measured using colorimetric assays for triglycerides (Infinity; Thermo Scientific), non-esterified fatty acids (Wako Diagnostics), and cholesterol (Cholesterol E, Wako Diagnostics). Plasma alanine aminotransferase (ALT) (MAK052, Sigma) and aspartate aminotransferase (AST) (MAK055, Sigma) were measured using dedicated kits according to manufacturer’s protocol.

### 2.4. Plasma lipoprotein analysis by fast protein liquid chromatography (FPLC)

Plasma lipoproteins were analyzed by running 200 μL of pooled plasma onto an FPLC system consisting of 2 Superose 6 columns connected in series (Amersham Pharmacia Biotech) as described previously [29].

### 2.5. Histology

#### 2.5.1. Hematoxylin & eosin staining

This was performed on formalin fixed liver, perigonadal white adipose and brown adipose tissues. Tissues were processed using ethanol and xylene before being embedded into paraffin blocks. Liver sections of 5 µm thickness were made using a microtome onto superfrost glass slides and incubated at 37 °C overnight. Tissue slices were then stained for 10 min in Mayer hematoxylin solution and for 10 seconds in eosin Y solution. The slides were then observed under a light microscope and representative images were taken.

### 2.5.2. Sirius red staining

Formalin fixed liver tissues embedded in paraffin blocks and sectioned at 5 µm thickness were deparaffinized in xylene, rinsed with deionized water and stained in solutions of phosphomolybdic acid for 2 minutes, picrosirius red F3BA stain for 60 minutes and 0.1N hydrochloride acid for 2 minutes. The staining was rinsed in 75% ethanol, dehydrated, air dried, and mounted with Depax mounting medium. Pictures were taken with light microscope.

#### 2.5.3. Trichome staining

This was performed on formalin-fixed 5 µm liver sections. The sections were deparaffinized, rehydrated and fixed for 1 hour in Bouin’s solution at 56°C. The slides were stained with Weigert’s iron hematoxylin, Biebrich scarlet-acid fuchsin, phosphomolybdic-phosphotungstic acid and aniline blue solutions. The slides were washed in distilled water, dehydrated, and mounted with resinous mounting medium before pictures were taken with light microscope.

### 2.6. Liver and fecal lipid extraction and quantification

Lipid extraction from liver and feces were performed as described (11). Liver and fecal lipids were measured using colorimetric assays for triglycerides (Infinity; Thermo Scientific) and cholesterol (Cholesterol E; Wako), according to manufacturer’s instruction and values were normalized by liver and fecal weight, respectively.

### 2.7. Fecal bomb calorimetry

Feces were collected from mice caged individually within a 24-h period during ad libitum feeding. Samples were dried for 48 hours at 60°C. Duplicate or triplicate samples of 45-55 mg were analyzed in Parr Instruments 6725 semimicro calorimeter using a 1109A Semimicro Oxygen Bomb placed in 450 grams of water. Heat of combustion was generated in kCal/g.

### 2.8. RNA Isolation and Quantitative Real-Time PCR

Total RNA was isolated from mouse tissues using TRIzol reagent (Life Technologies). 2000 ng of RNA was used to synthesize cDNA by reverse transcription using High-Capacity cDNA Reverse Transcription Kit (Applied Biosystems). Gene expression was performed by quantitative PCR with iTaq Universal SYBR Green Supermix (Bio-Rad). Gene expression was normalized to *36b4* as a housekeeping gene. Primer sequences for measured genes are listed in Supplemental Table 1.

### 2.9. Statistical Analysis

Data are presented as Means ± Standard Error. Statistical analysis by Student’s t-test, one-way ANOVA or 2-way ANOVA followed by a Tukey’s post hoc multiple comparison test. p < 0.05 is considered statistically significant.

## 3. RESULTS

### 3.1. Knockout of Cyp2c genes results in a human-like BA profile and a reduction in 12α-hydroxylated BAs

Heterozygous Cyp2c^+/−^ mice were bred to generate Cyp2c^−/−^, Cyp2c^+/−^, and WT control mice (Figure 1A). Cyp2c44 is a member of this family of Cyp2c genes but is physically located outside the cluster and was therefore intact (Figure 1A). In the WT liver tissues, *Cyp2c29, Cyp2c37, Cyp2c38, Cyp2c40, Cyp2c50, Cyp2c54, Cyp2c69* and *Cyp2c70* were highly expressed with cycle threshold (Ct) values of 22 or lower by qPCR. *Cyp2c67, Cyp2c39, Cyp2c55* and *Cyp2c68* were moderately expressed, whereas *Cyp2c65* and *Cyp2c66* were lowly expressed with Ct values ~35 in the WT mice. All of these mRNAs were modestly reduced in the Cyp2c^+/−^ mice and undetectable in the Cyp2c^−/−^ mice (Figure 1B). As expected, mRNA expression of *Cyp2c44* –which is located outside of the deleted gene cluster– did not differ between the 3 genotypes (Figure 1B).

**Figure 1.**
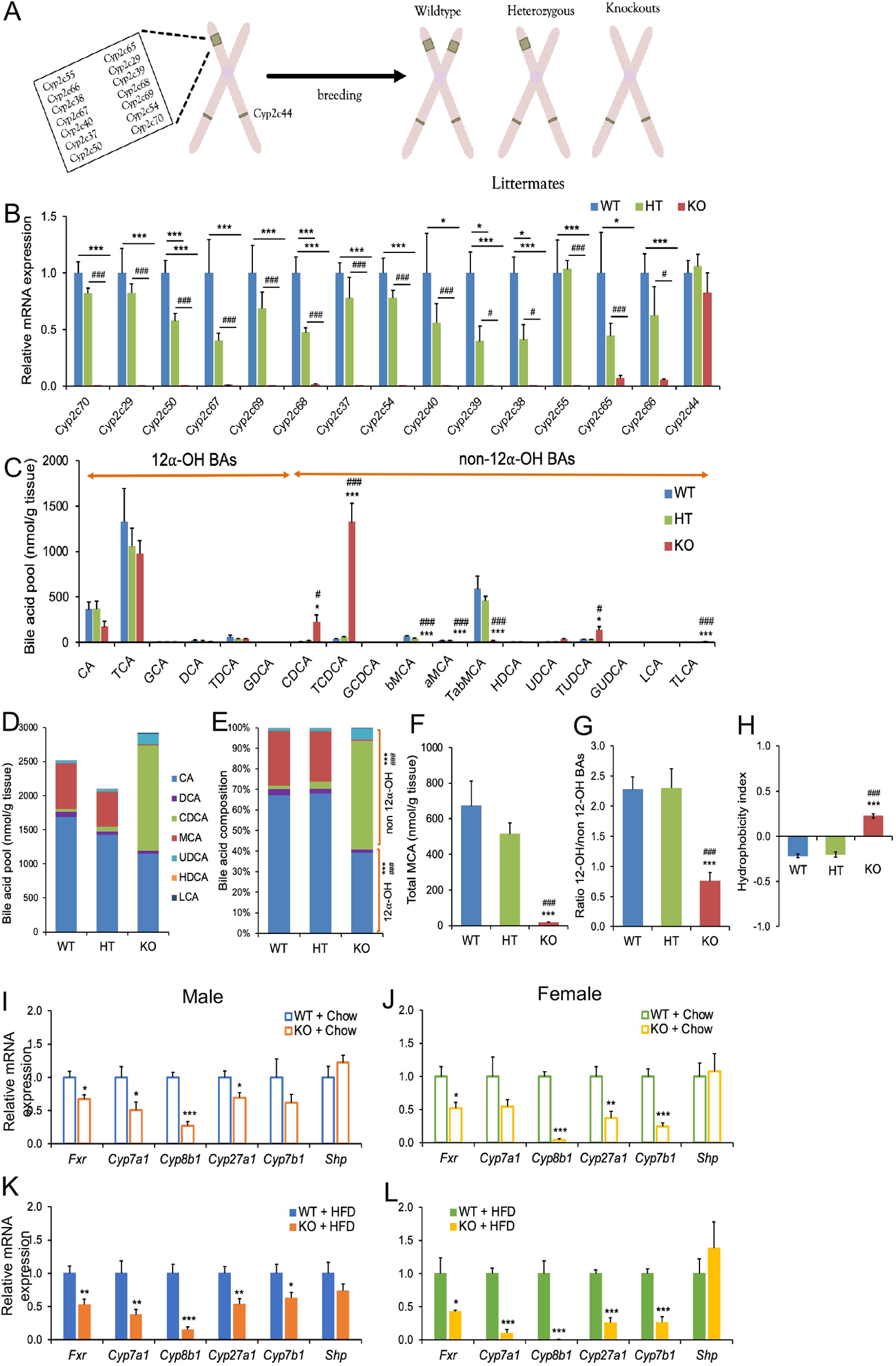
Cyp2c deficiency humanizes murine BA pool due to MCA depletion. A: Schematic of the 14 Cyp2c genes that were deleted in Cyp2c^−/−^ model and the progeny genotypes from breeding heterozygotes. B: mRNA expression of Cyp2c genes in WT, Cyp2c^+/−^ and Cyp2c^−/−^ mice. C: BA species measured by UPLC-MS/MS from a pool of liver, gallbladder, and intestine. D-E: The absolute quantities (D) and relative proportions (E) of BAs. For D-F, each BA represents a total of unconjugated, taurine-conjugated, and glycine-conjugated subspecies. F: Total levels of all MCAs in WT, Cyp2c^+/−^ and Cyp2c^−/−^ mice. G: Ratio of total 12α-OH to total non-12α-OH BAs. H: Hydrophobicity index of BAs. I-L: Hepatic mRNA expression of BA metabolism genes in males on chow diet (I), females on chow diet (J), males on HFD (K), and females on HFD (L). Data are represented as mean ± SEM. n=6 mice/group for Figures B-H, 5-7 mice/group for Figures I-L. *p< 0.05, **p< 0.01 and ***p<0.001 relative to WT and #p< 0.05, and ###p<0.001 relative to Cyp2c^+/−^. B-H by one-way ANOVA; I-L by Student’s t-test.

UPLC-MS/MS analysis of BAs was performed on a pool of liver, gallbladder and small intestine in males and females from each of the 3 genotypes. There were no sex-dependent differences in BA composition, so the male and female data were pooled together. The reduced expression of a number of Cyp2c genes in the Cyp2c^+/−^ mice did not result in any significant changes in BA levels or composition compared to the WT mice, whereas the Cyp2c^−/−^ mice showed significant changes in the BA composition compared to Cyp2c^+/−^ and WT mice (Table 1 and Figures 1C, D). The levels of conjugated and unconjugated MCA were hardly detectable in the Cyp2c^−/−^ mice, validating that deletion of the Cyp2c cluster generates an MCA-deficient human-like BA profile (Figures 1C-F). On the other hand, there was an increase in the levels of conjugated and unconjugated CDCA in the Cyp2c^−/−^ mice, consistent with CDCA being the predominant precursor for MCA synthesis (Figures 1D, E, F). Tauroursodeoxycholic acid (TUDCA) and taurolithocholic acid (TLCA)–which can be derived from CDCA–were increased in the Cyp2c^−/−^ mice (Figures 1C, D, E). The absence of MCAs and compensatory increase in CDCA demonstrate that Cyp2c^−/−^ mice have a human-like BA pool.

**Table 1.**
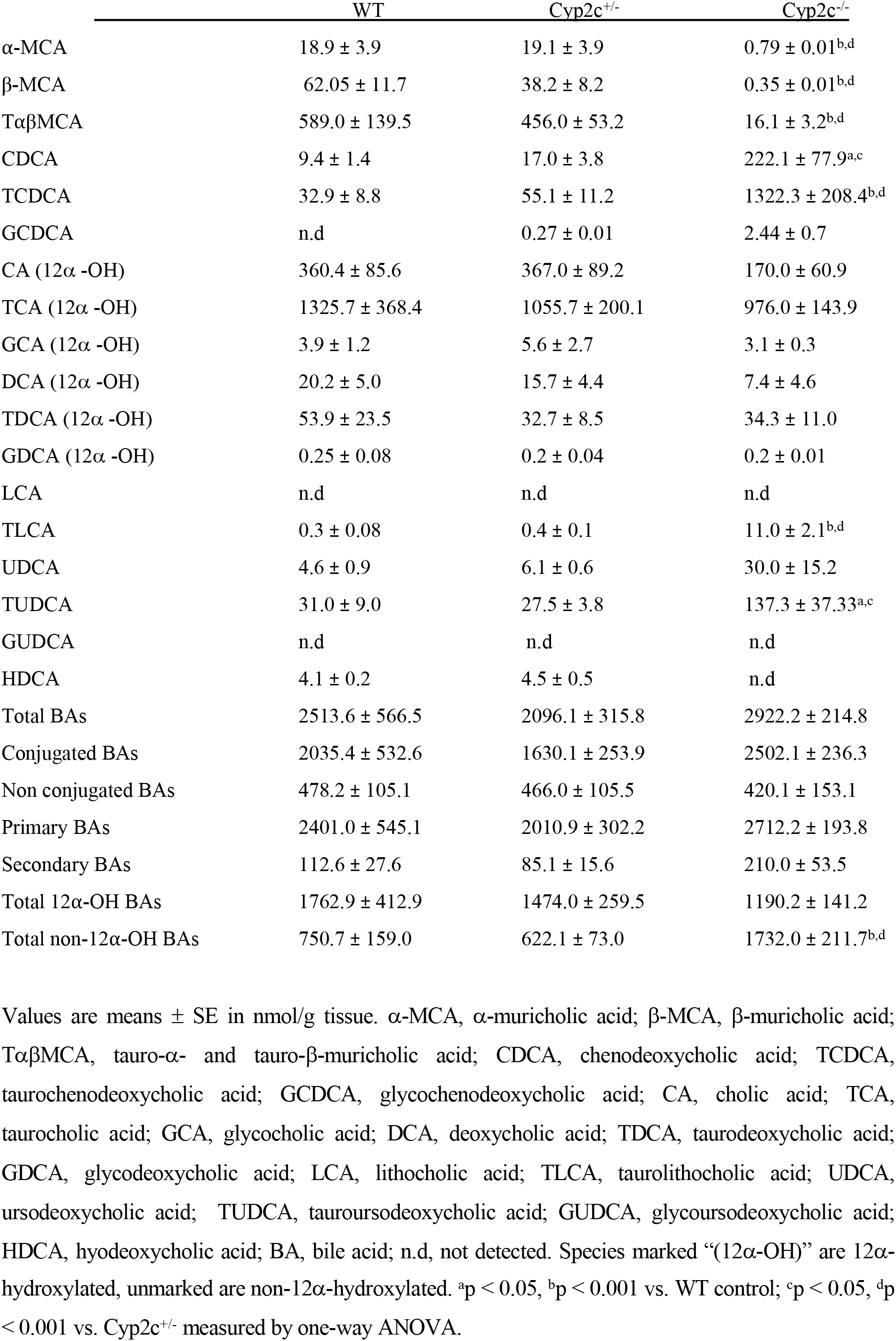
Pooled bile acid profile by UPLC-MS/MS

We further characterized the BA pool composition based on other important physicochemical BA features. There were no differences between genotypes in the total amounts of conjugated or unconjugated BAs, and no differences in the total amounts of primary or secondary BAs (Table 1). In the Cyp2c^−/−^ mice, a smaller portion of the BA pool was made up of BAs that are hydroxylated at the 12α-carbon position (12α-OH BAs)–cholic acid (CA), its secondary BA derivative deoxycholic acid (DCA), and their conjugated forms (Figures 1E). These BAs are generated by the enzymatic activity of Cyp8b1, the 12a-hydroxylase. On the other hand, non-12α-OH BAs were significantly increased, leading to a significant decrease in the ratio of 12α-OH/non-12α-OH BAs (Figure 1G and Table 1). This is consistent with what was observed in Cyp2c70-deficient mice [20,30]. MCA is considered a hydrophilic BA, therefore its depletion and resultant alteration in BA composition increased the hydrophobicity index of BA pool in the Cyp2c^−/−^ mice as calculated according to Heuman [31] (Figure 1H). Because there were no differences in BA profile between WT and Cyp2c^+/−^ mice, subsequent experiments were conducted only in WT and Cyp2c^−/−^ mice. Moreover, we conclude that partial reductions in expression of Cyp2c genes are insufficient to modulate BA pool composition.

To test whether changes in expression of BA metabolism genes could explain the altered BA composition in Cyp2c^−/−^ mice, we measured the expression of key BA metabolism genes. In males and females on chow or HFD, there was a reduction of key BA synthesis genes *Cyp7a1, Cyp8b1, Cyp27a1* and *Cyp7b1*, along with decreased expression of *Fxr* in the liver of the Cyp2c^−/−^ mice (Figures 1I-L). There were no significant changes in hepatic expression of *Shp* (Figure 1I-L), and there were no differences in ileal expression of *Fxr* or *Fgf15* between WT and Cyp2c^−/−^ mice (data not shown). The reduction in *Cyp8b1* expression is consistent with the smaller proportion of 12α-OH BAs in Cyp2c^−/−^ mice.

These changes in BA composition suggest that Cyp2c^−/−^ mice may have alterations in intestinal lipid absorption that could affect susceptibility to HFD-induced obesity. The effects could be predicted to go in two opposing directions. Eliminating 12α-OH BAs from WT mice is known to reduce fat absorption [13,14,16]. Thus the relative reduction in 12α-OH BAs in Cyp2c^−/−^ mice suggests that they may be less susceptible to HFD-induced obesity. On the other hand, the increase in hydrophobicity of the BA pool when MCAs are absent might be predicted to enhance intestinal lipid absorption and promote obesity. We next investigated how the changes in BA composition affect obesity susceptibility and energy metabolism in the Cyp2c^−/−^ mice.

### 3.2. Male Cyp2c^−/−^ mice are resistant to HFD-induced obesity

In mice on chow diet, weekly body weight measurements showed no significant differences between WT and Cyp2c^−/−^ mice in males (Figure 2A) or females (Figure 2B), and there were no differences in average food intake between the WT and Cyp2c^−/−^ mice in either sex (Figure 2C, D). Body composition analysis showed no differences in lean and fat mass between the genotypes in chow-fed males, but chow-fed female Cyp2c^−/−^ mice had slightly higher lean mass than WT controls (Figure 2E, F).

**Figure 2.**
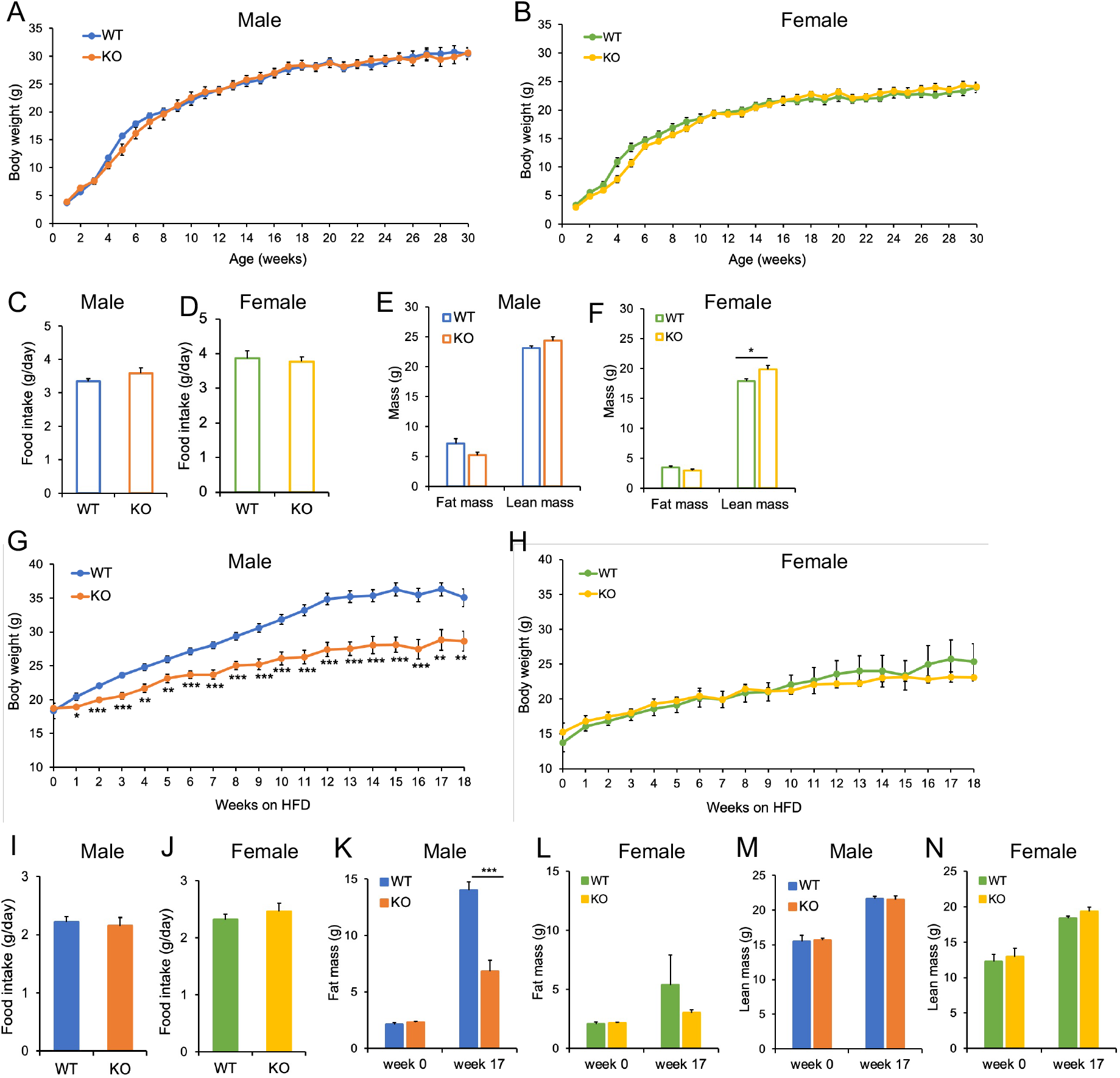
Cyp2c deficiency protects against diet induced obesity in male mice. A-B: Weekly body weight in male (A) and female (B) mice on chow diet. C-D: Average daily food intake in male (C) and female (D) chow-fed mice. E-F: Fat and lean mass in male (E) and female (F) chow-fed mice. G-H: Weekly body weight in male (G) and female (H) fed HFD. I-J: Average daily food intake in male and female (J) mice fed HFD. K-L: Fat mass in males (K) and females (L) before and after 17 weeks on HFD. M-N: Lean mass in males (M) and females (N) before and after 17 weeks on HFD. Data are represented as mean ± SEM. n=7 mice/group for all groups, except n=5 mice/group for females + HFD. *p< 0.05, **p< 0.01 and ***p<0.001 relative to WT by Student’s t-test.

In a parallel study conducted in a second cohort of mice fed a HFD (60% of kcal derived from fat) for 18 weeks, body weights of males diverged after just one week, such that male WT mice gained substantial body weight, whereas male Cyp2c^−/−^ mice were protected (Figure 2G). Female mice of both genotypes were protected from obesity (Figure 2H). We did not detect any significant difference in average food intake between genotypes (Figure 2I, J). Prior to HFD intervention, fat mass was similar between genotypes, but after 17 weeks on HFD, fat mass was significantly lower in Cyp2c^−/−^ males compared to WT controls (Figure 2K). We did not observe any significant differences between genotypes in females (Figure 2L). Lean mass did not differ between the genotypes (Figure 2M, N). This demonstrates that the low body weight of male Cyp2c^−/−^ mice was due to low adiposity.

### 3.3. Male and female Cyp2c^−/−^ mice show improvements in glucose homeostasis

Cyp2c^−/−^ male mice showed significant improvements in OGTT on both chow and HFD (Figure 3A, B) as well as improvements in ITT on chow diet but not on HFD (Figure 3C, D). Cyp2c^−/−^ females on both chow and HFD showed lower glucose 15-minutes after glucose gavage during OGTT, but no significant difference in total area under the curve (Figure 3E, F). Cyp2c^−/−^ females also showed improved ITT on chow diet but not on HFD (Figure 3G, H).

**Figure 3.**
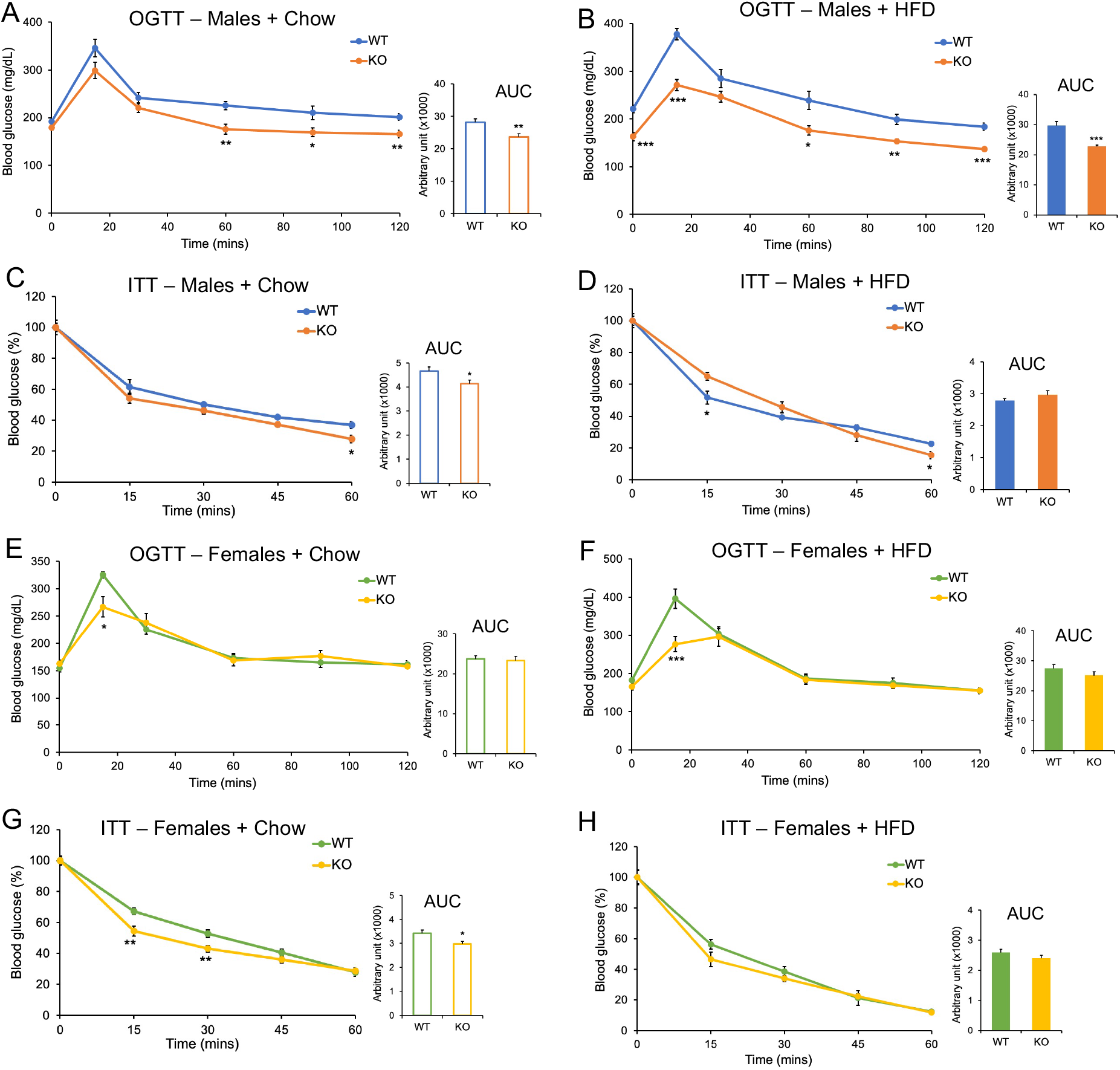
Cyp2c deficiency improves systemic glucose homeostasis independent of diet and sex. A-B: Blood glucose level during OGTT in males under chow (A) or HFD (B) conditions. C-D: Blood glucose level during ITT in males under chow (C) or HFD (D) conditions. E-F: Blood glucose level during OGTT in females under chow (E) or HFD (F) conditions. G-H: Blood glucose level during ITT in females under chow (G) or HFD (H) conditions. Insets represent area under the curve (AUC). Data are represented as mean ± SEM. n=7 mice/group for all groups, except n=5 mice/group for females + HFD. *p< 0.05, **p< 0.01 and ***p<0.001 relative to WT by Student’s t-test.

To assess whether these improvements in OGTT were specific to an incretin effect, we also performed IPGTTs. Male Cyp2c^−/−^ mice showed improved IPGTTs on both diets compared to WT (Supplemental Figure 1A, B), and males of all diets and genotypes showed a strong incretin effect (Supplemental Figure 1C-F). In females, there were no differences in IPGTT between genotypes (Supplemental Figure 1G, H), and no incretin effect on either diet (Supplemental Figure 1I-L).

We further analyzed plasma samples obtained from mice that were fasted for 5-hours. Under chow diet conditions, there were no differences in blood glucose or plasma insulin between genotypes, but under HFD conditions, Cyp2c^−/−^ male but not Cyp2c^−/−^ female mice showed significantly lower levels of blood glucose and plasma insulin compared to WT controls (Figure 4A, B). Taken together, these data suggest that Cyp2c^−/−^ mice of both sexes have improvements in systemic glucose homeostasis that are independent of incretin hormones.

**Figure 4.**
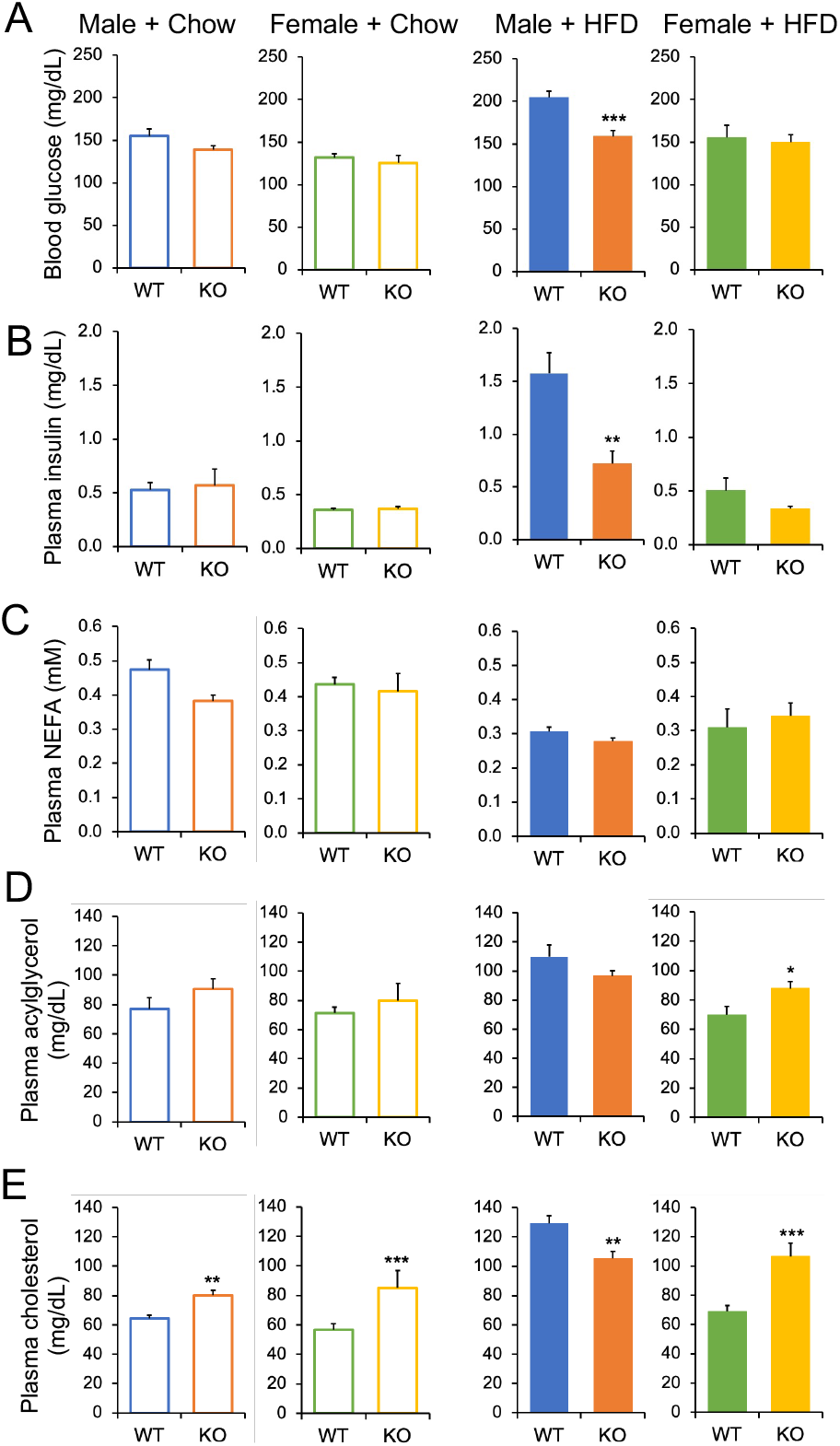
Effects of Cyp2c deficiency on plasma metabolites and insulin. A-E: Blood glucose (A) and plasma levels of insulin (B), NEFA (C), acylglycerol (D), and cholesterol (E) levels in males and females fed chow or HFD, after a 5-hour fast. Data are represented as mean ± SEM. n=7 mice/group for all groups, except n=5 mice/group for females + HFD. *p< 0.05, **p< 0.01 and ***p<0.001 relative to WT by Student’s t-test.

### 3.4. Cyp2c-deficiency alters plasma lipoprotein profile

After a 5-hour fast, plasma levels of non-esterified fatty acids (NEFA) did not differ between the mice (Figures 4C). On chow diet, plasma acylglycerols did not differ between genotypes, but on HFD plasma acylglycerols were significantly increased in the Cyp2c^−/−^ females compared to WT controls (Figure 4D). Plasma cholesterol was increased in Cyp2c^−/−^ males and Cyp2c^−/−^ females on chow diet, as well as in Cyp2c^−/−^ females on HFD, showing that the lack of MCAs generally raises circulating cholesterol [20,32]. On the other hand, under HFD conditions, WT males had high cholesterol levels, and this was significantly blunted in the Cyp2c^−/−^ males (Figures 4I, J), possibly as a secondary consequence to their obesity resistance.

We next examined lipoprotein subfractions. In general, we found that Cyp2c^−/−^ mice had higher levels of cholesterol and acylglycerol in LDL fractions, and this effect was stronger in females (Figure 5A-H). To test for potential explanations for the changes in the lipoprotein profile, we measured mRNA expression of key cholesterol metabolism genes. While the transcription factor Srebp2 and several of its target genes were not different between genotypes, we found that LDL-receptor (*Ldlr*) expression tended to be decreased in Cyp2c^−/−^ mice (Figure 5I-L). These data suggest that lack of MCAs may lower the clearance of LDL particles via reductions in hepatic *Ldlr*, as previously suggested [20,32].

**Figure 5.**
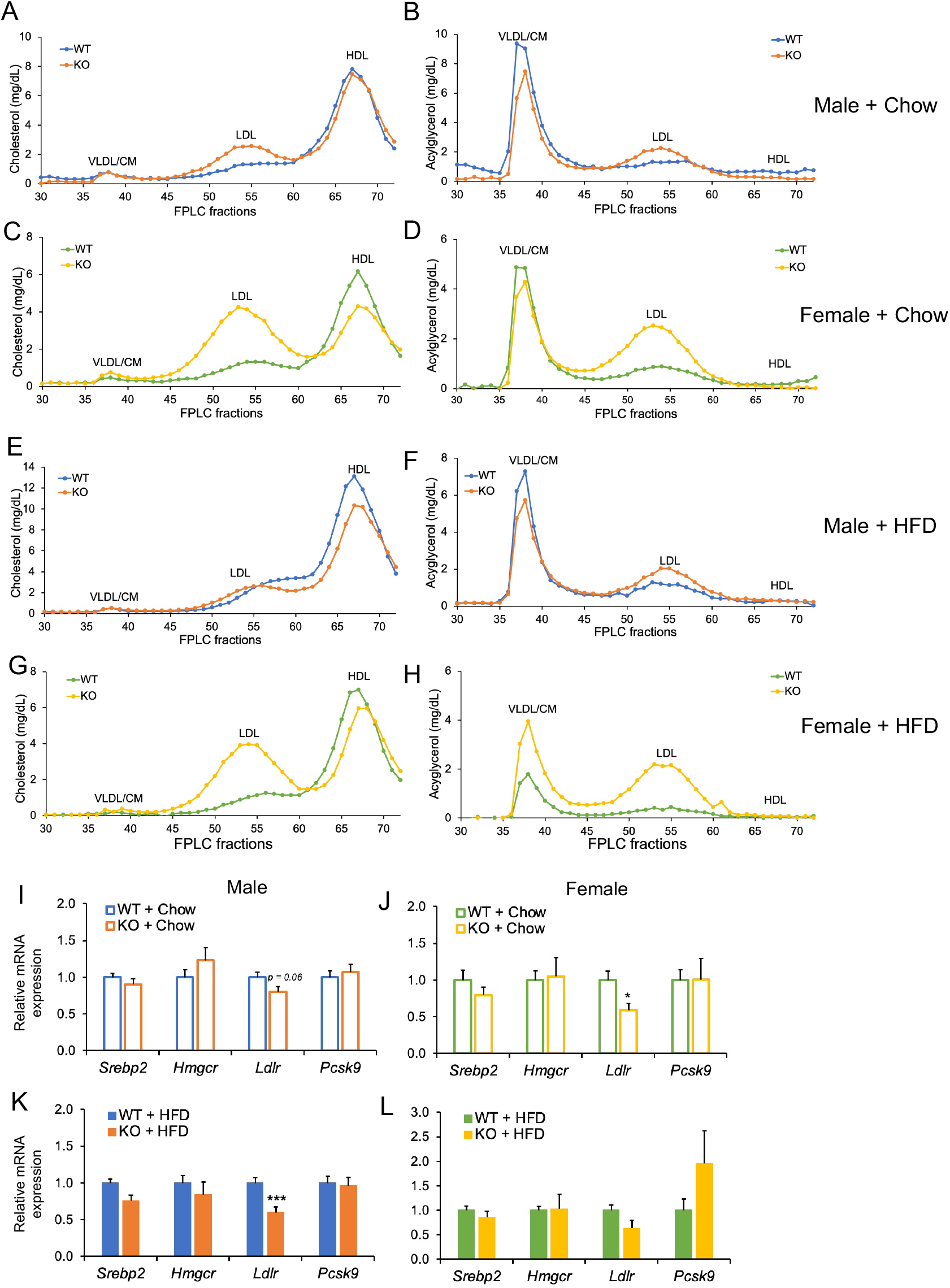
Cyp2c deficiency alters lipoprotein profile. A-B: Plasma cholesterol (A) and acylglycerol (B) in FPLC fractions of male chow-fed mice. C-D: Plasma cholesterol (C) and acylglycerol (D) in FPLC fractions of female chow-fed mice. E-F: Plasma cholesterol (E) and acylglycerol (F) in FPLC fractions of male HFD-fed mice. G-H: Plasma cholesterol (G) and acylglycerol (H) in FPLC fractions of female HFD-fed mice. I-L: Hepatic mRNA expression of cholesterol metabolism genes in males on chow diet (I), females on chow diet (J), males on HFD (K), and females on HFD (L). Data are represented as mean ± SEM. n=7 mice/group for all groups, except n=5 mice/group for females + HFD. *p< 0.05, **p< 0.01 and ***p<0.001 relative to WT by Student’s t-test.

### 3.5. Resistance of Cyp2c^−/−^ male mice to HFD-induced obesity protects against adipose inflammation

Cyp2c^−/−^ mice tended to have reductions in relative gonadal fat mass, in both sexes (Figure 6A). Chronic-low grade inflammation and infiltration of immune cells are well-known hallmarks of increased adiposity [33,34]. Consistent with this, we observed crown-like structures and larger adipocytes in the gonadal fat of the HFD-fed male WT mice (Figure 6B) but not in other male groups or female groups. In line with the histology, the obesity-sensitive male WT mice showed a strong increase in expression of inflammatory genes such as *F4/80, Cd68, Mcp1* and *Tnfa*, but not in the other male groups or female groups (Figure 6C, D). It has been reported that in obesity, de-novo lipogenesis in the fat tissue is repressed [35,36]. In agreement, lipogenic genes such as *Srebp1c* and its downstream targets trended towards decreased expression in the HFD-fed male WT mice, but this effect was blunted in the non-obese male and female groups (Figure 6E, F). Taken together, Cyp2c^−/−^ male mice show adipose tissue phenotypes consistent with their decreased obesity.

**Figure 6.**
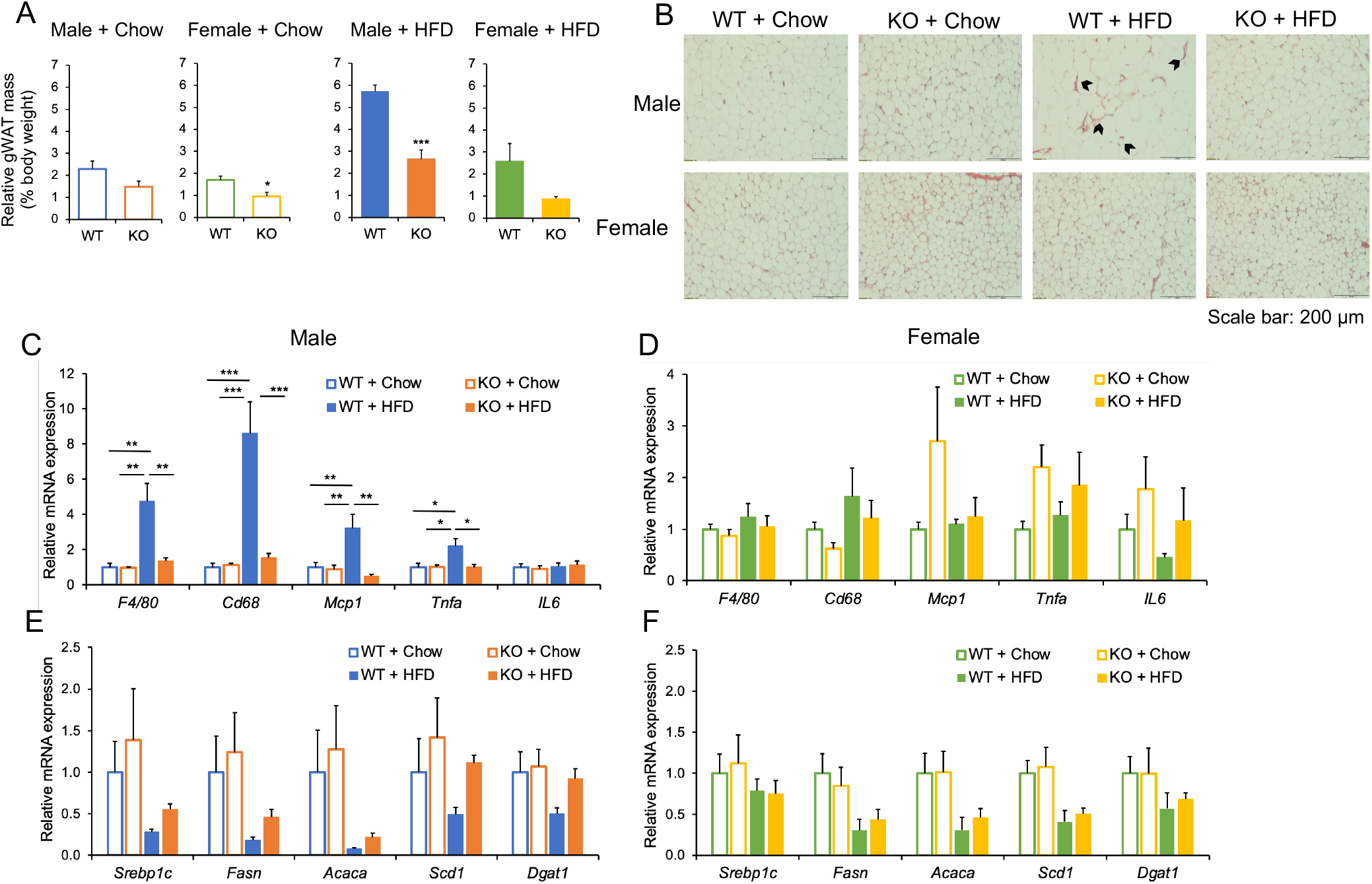
Resistance to obesity in Cyp2c^−/−^ mice protects against adipose tissue inflammation. A: Gonadal white adipose tissue (gWAT) mass relative to body weight in males and females fed chow or HFD. B: Representative images of H & E staining of gWAT in males and females. Scale bar is 200 µm. C-D: mRNA expression of pro-inflammatory genes in gWAT of males (C) and females (D) on chow or HFD. E-F: mRNA expression of lipogenesis genes in gWAT of male (E) and female (F) on chow diet or HFD. Data are represented as mean ± SEM. n=7 mice/group for all groups, except n=5 mice/group for females + HFD. *p< 0.05, **p< 0.01 and ***p<0.001 by two-way ANOVA and Tukey’s multiple comparisons test.

### 3.6. Energy Balance

To examine the mechanisms of reduced diet-induced obesity in the male Cyp2c^−/−^ mice, we performed indirect calorimetry in male mice. We detected no significant differences in energy expenditure between the genotypes on chow diet (Figures 7A) or HFD (Figures 7B). Additionally, histological staining of brown adipose tissue (Supplemental Figure 2A) and expression of thermogenic genes showed no differences between the groups on either chow or HFD (Supplemental Figure 2B, C).

**Figure 7.**
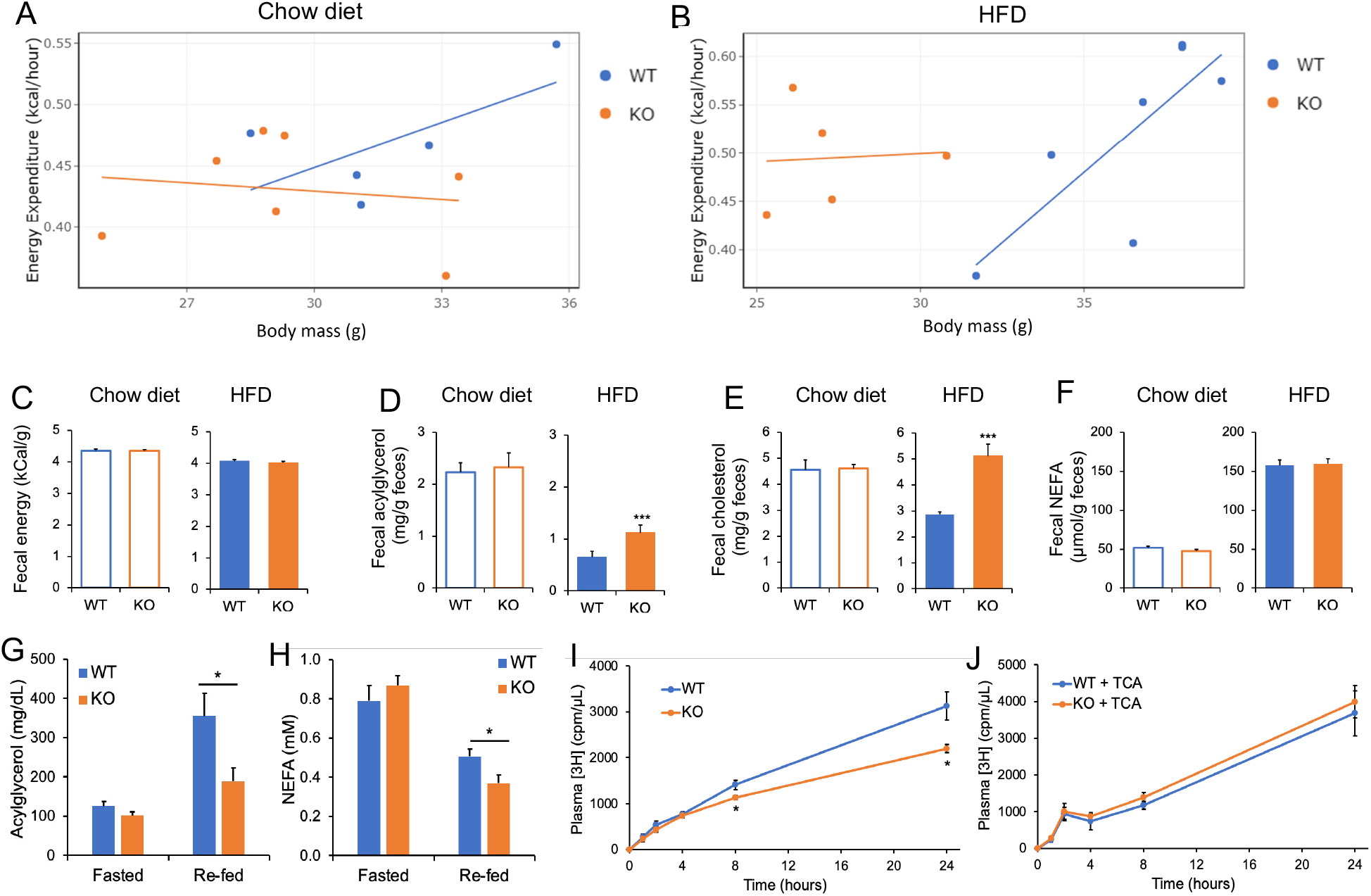
Indirect calorimetry and lipid absorption analyses in male Cyp2c^−/−^ mice. A-B: Average hourly energy expenditure via indirect calorimetry over a 4-day period for male mice on chow (A) and HFD (B). C: Fecal energy content by bomb calorimetry in mice on chow and HFD diets. D-F: Fecal levels of acylglycerol (D), cholesterol (E) and NEFA (F) in chow-fed and HFD-fed mice. G-H: Plasma levels of acylglycerol (G) and NEFA (H) in mice fasted overnight for 14 hours and re-fed HFD for 2 hours. I-J: Plasma levels of [^3^H] in mice undergoing the radiolabeled triolein absorption experiment without (I) or with (J) 3 consecutive days of low-dose TCA treatment. Data are represented as mean ± SEM. n=5-7 mice/group for Figures A-F, 5-6 mice/group for Figures G-J. *p< 0.05, **p< 0.01 and ***p<0.001 relative to WT by Student’s t-test.

Because we observed substantial protection from HFD-induced obesity in Cyp2c^−/−^ males, but no significant differences in food intake or energy expenditure, we reasoned that there may be an increase in calorie excretion. However, bomb calorimetry showed no significant differences in fecal energy output between the Cyp2c^−/−^ mice and WT controls on either chow or HFD (Figure 7C). Our interpretation of these unexpected findings is that male Cyp2c-/-mice on a HFD have slight changes in food intake, energy expenditure, calorie excretion, or all three, that are below the limit of detection in these assays, but that add up over time to significant protection from obesity.

### 3.7. Cyp2c^−/−^ mice have impaired intestinal lipid absorption, which can be rescued by taurocholic acid

A key premise of this study is that MCAs are thought to promote poor lipid absorption, and therefore eliminating MCAs and shifting the mouse BA pool to other types of BAs may be predicted to enhance lipid absorption. To pursue this, we quantified fecal lipids. During chow feeding, there were no changes in fecal acylglycerol, cholesterol or NEFA between the two genotypes (Figures 7D-F). However, during HFD feeding, fecal acylglycerol and cholesterol content, but not NEFA, were significantly increased in the stool of Cyp2c^−/−^ male mice (Figures 7D-F), suggesting that under HFD conditions, Cyp2c^−/−^ may have increased lipid excretion and reduced lipid absorption compared to WT controls.

To further assess whether Cyp2c^−/−^ mice have reduced intestinal lipid absorption, we performed two additional experiments. First, we performed a fasting re-feeding experiment. Following an overnight fast (~14 hours), plasma levels of acylglycerol (Figure 7G) and NEFA (Figure 7H) did not differ between WT and Cyp2c^−/−^ mice. After 2 hours of re-feeding HFD, plasma acylglycerol and NEFA were significantly lower in the Cyp2c^−/−^ mice (Figure 7G, H), consistent with reduced intestinal lipid uptake.

In the second experiment, we used a radiolabeled triolein to directly measure intestinal lipid absorption. We fasted mice for 4-hours, injected them with poloxamer 407 to block lipoprotein clearance, then administered olive oil containing a radiolabeled triglyceride, [^3^H] triolein, by oral gavage. The Cyp2c^−/−^ mice showed significantly lower plasma levels of [^3^H] at 8- and 24-hours post-gavage, demonstrating reduced absorption of intestinal triglyceride compared to WT (Figure 7I).

We reasoned that the impaired triglyceride absorption in the Cyp2c^−/−^ mice could be due to their reduction in the ratio of 12α-OH/non-12α-OH BAs. To test whether increasing 12α-OH BAs in Cyp2c^−/−^ mice rescues their defect in intestinal lipid absorption, WT and Cyp2c^−/−^ mice received an oral gavage of either a low dose (17 mg/kg) of taurocholic acid (TCA) or vehicle for 3 consecutive days. The 17 mg/kg dose was used because of previous reports that this dose elevates levels of CAs without changing the total BA levels [26,27]. The TCA gavage rescued the lipid absorption of Cyp2c^−/−^ mice, as demonstrated by the lack of any differences in radioactive tracers in the plasma of the Cyp2c^−/−^ mice compared to WT (Figure 7J). Taken together, Cyp2c deficiency and the resultant relative reduction in 12α-OH BAs promotes fecal lipid excretion and reduces intestinal lipid absorption.

### 3.8. Cyp2c^−/−^ mice have liver injury

The liver is central to lipid and BA metabolism and therefore responsive to modulated BA metabolism. Compared to WT controls, liver mass relative to body weight was increased in Cyp2c^−/−^ mice (Figure 8A). This was not due to accumulation of excess hepatic lipid, as acylglycerols were significantly reduced in Cyp2c^−/−^ mice (Figures 8B). Liver cholesterol did not differ between genotypes in male mice, but female Cyp2c^−/−^ mice showed a significant reduction in liver cholesterol under HFD condition (Figure 8C).

**Figure 8.**
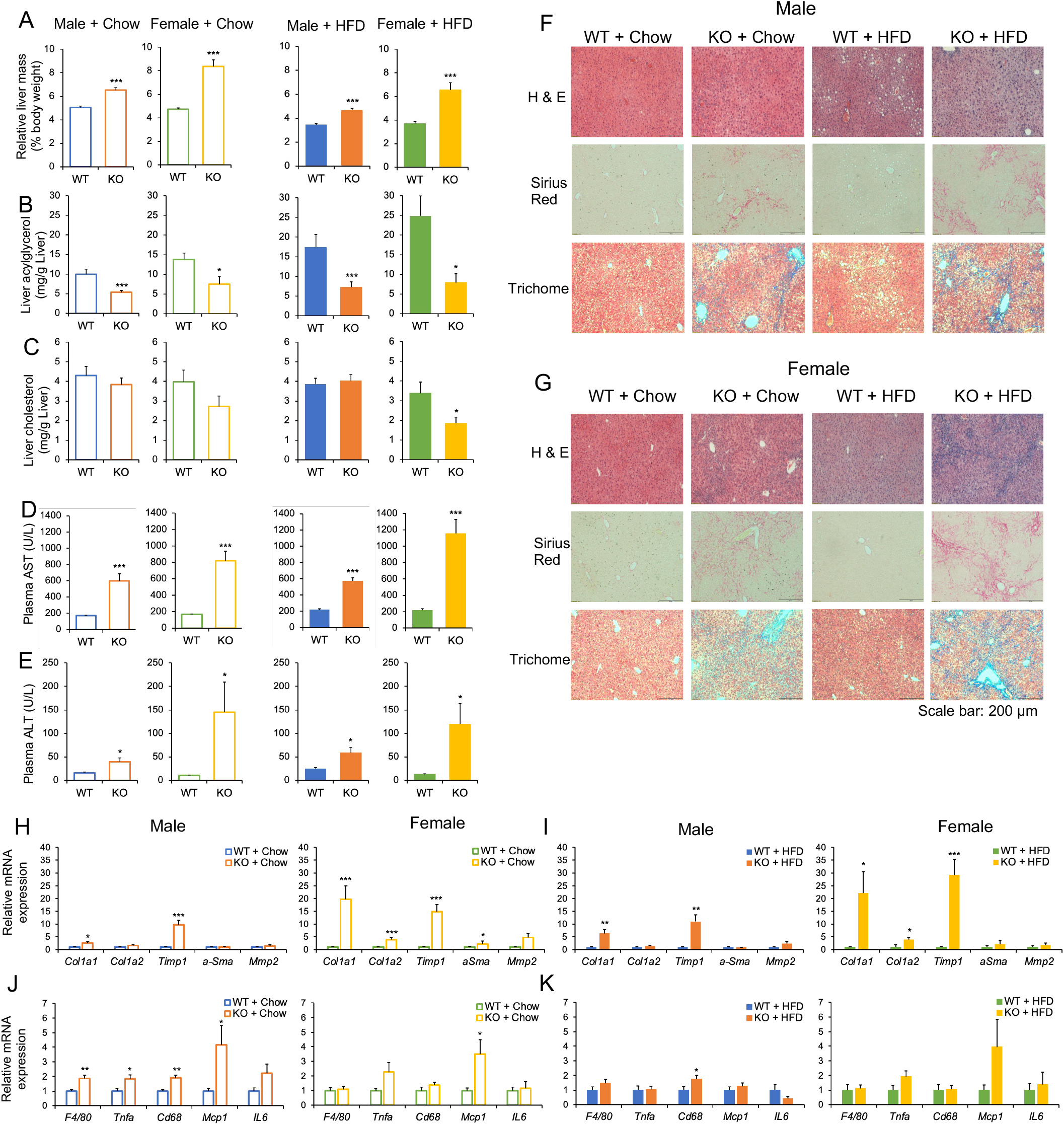
Cyp2c^−/−^ mice have liver injury. A: Liver mass relative to body weight in males and female on chow or HFD. B: Hepatic acylglycerol in male and female mice on chow or HFD. C: Hepatic cholesterol in male and female mice on chow or HFD. D: Plasma AST levels in male and female mice on chow or HFD. E: Plasma ALT levels in male and female mice on chow or HFD. F-G: Representative images from H & E, Sirius red, and trichome stainings in males (F) and females (G) fed chow or HFD. Scale bar is 200µm. H-I: Hepatic mRNA expression of fibrosis genes in male and female mice on chow (H), and male and female on HFD (I). J-K: Hepatic mRNA expression of pro-inflammatory genes in male and female mice on chow (J), and male and female on HFD (K). Data are represented as mean ± SEM. n=7 mice/group for all groups, except n=5 mice/group for females + HFD. *p< 0.05, **p< 0.01 and ***p<0.001 relative to WT by Student’s t-test.

Increased hepatotoxicity has been reported in mice fed diets supplemented with LCA [37,38]. In rat hepatocytes, CDCA induced hepatotoxicity more potently than other BAs [39]. Because CDCA and TLCA levels are elevated in Cyp2c^−/−^ mice, we assessed liver damage. Plasma levels of liver enzymes AST and ALT were significantly increased in both male and female Cyp2c^−/−^ mice in a diet-independent manner, with the females showing stronger elevations compared to males (Figures 8D, E). In male mice, Cyp2c deficiency resulted in mild liver fibrosis, as evidenced by increased collagen deposition in Sirius red and trichome stains (Figure 8F). In female mice, Cyp2c deficiency caused strong liver fibrosis (Figure 8G). In agreement with the histological data, mRNA expression of fibrosis gene markers corroborated the mild fibrosis in the Cyp2c^−/−^ male mice and the strong fibrosis in the Cyp2c^−/−^ female mice (Figure 8H-I). Additionally, there was a modest increase in inflammatory gene markers in the Cyp2c^−/−^ male and female mice (Figures 8J, K). Taken together, Cyp2c deficiency leads to hepatic injury, which is more severe in females than males.

## 4. DISCUSSION

In this study, we confirm the depletion of MCAs in Cyp2c-deficient mice, and further show that mice that are heterozygous for Cyp2c retain their ability to synthesize normal levels of MCAs, despite 50% decreases in the expression of most Cyp2c genes. Furthermore, we show that contrary to predictions, MCA *depletion* results in impaired lipid absorption and protection against high fat diet-induced obesity in male mice. Additionally, Cyp2c deficiency promotes mild and severe liver injury in male and female mice, respectively, consistent with a recent publication [30].

Published studies have suggested that mice with high levels of MCAs are protected against high fat diet-induced obesity [15,16], potentially because MCAs are considered hydrophilic BAs that are predicted to poorly promote intestinal lipid absorption. However, those studies indirectly raised MCAs, as a secondary consequence of Cyp8b1 deletion. Our study directly investigates the role of MCAs in response to obesogenic diet, by using mice that lack MCAs. Our results disagree with the prediction that MCAs protect against obesity. We found that in both males and females, the absence of MCA did not promote obesity. In fact, in males, absence of MCA *protected* against high fat diet-induced obesity and the associated chronic low-grade inflammation of the adipose tissue. We tested whether this protection was explained by increased energy expenditure, decreased food intake, or increased calorie excretion, but did not find a significant difference in any single parameter that could explain the low body weight. Our interpretation is that mild differences in some or all of these parameters likely exist but are not detectable with our instruments and statistical power. Of note, absence of MCA disrupted the relationship between energy expenditure and body mass. In wild type C57Bl/6 mice, larger mice require greater levels of energy expenditure [40]. However, this correlation is not observed in Cyp2c deficient mice (Fig 7A, 7B). A similar effect was observed in mice engineered with a microbiome lacking a bile salt hydrolase [41], suggesting a possible role for the microbiome that will be of interest for future studies.

We also directly tested the expectation, based on prior publications, that lack of MCAs would raise the hydrophobicity of the BA pool and consequently increase intestinal lipid absorption. Our results showed the opposite effect–despite a significant increase in BA hydrophobicity index in the knockout mice, there was a significant decrease in intestinal triglyceride absorption. We conclude that the hydrophobicity index of a BA pool is insufficient to explain functional effects of BAs on intestinal lipid absorption.

On the other hand, in Cyp2c^−/−^ mice, there is a significant decrease in the ratio of 12α-OH/non-12α-OH BAs. This phenotype is likely due to the strong repression of Cyp8b1, the enzyme responsible for synthesizing 12α-OH BAs, as previously suggested [20,32]. Indeed, studies in mice from our group and others have shown that knockout of Cyp8b1 depletes the murine BA pool of 12α-OH BAs, and this leads to reduced intestinal lipid absorption, lower body weight, and improvements in cardiometabolic parameters [14–17]. Thus, the decreased lipid absorption of Cyp2c^−/−^ mice is consistent with the reduced ratio of 12α-OH/non-12α-OH BAs in these mice. This is further supported by our finding that an oral gavage of TCA, the major 12α-OH BA, rescues the intestinal lipid absorption in the Cyp2c^−/−^ mice. Altogether, these data suggest that 12α-OH BA content of the BA pool is more important than hydrophobicity index for regulating the efficiency of intestinal lipid absorption.

Our work shows that Cyp2c deficiency also causes other interesting metabolic effects. For one thing, it causes a more human-like LDL-cholesterol-enriched lipoprotein profile, particularly in females. In our study as well as the studies in Cyp2c70-knockdown mice, observed downregulation of the LDL-receptor potentially contributes to the elevated LDL-cholesterol [30,32]. Another change we observed was the improvement in glucose metabolism. This improvement appears to be independent of lower body weight, as we observed in female Cyp2c^−/−^ mice and in chow-fed Cyp2c^−/−^ mice, which have no differences in body weight compared to controls. This effect is also somewhat contrary to what might be predicted based on the literature. Prior studies have shown that glycine and taurine conjugated β-MCA can antagonize intestinal FXR leading to improvements in glucose homeostasis and metabolic disorders [11,42]. Our data suggests that MCA *depletion* confers improved glucose homeostasis. Future studies will be required to determine the mechanisms of this improvement, but it appears to be independent of the intestine, based on our OGTT and IPGTT data.

Finally, our study shows liver damage in the Cyp2c^−/−^ mice, which appears more severe in females than males as also recently reported [30]. It has previously been observed that elevated levels of certain BAs including LCA and CDCA can be hepatotoxic [37,38]. Thus, the increases in CDCA and LCA may contribute to the liver damage phenotypes in the Cyp2c^−/−^ mice as others have suggested [30]. Because Cyp2c enzymes are also involved in drug metabolism and detoxifying xenobiotics [43,44], we cannot rule out that these roles also contribute to the effects of Cyp2c deletion on liver damage.

In conclusion, our study shows that the absence of MCA in Cyp2c-deficient mice results in altered BA composition characterized by decreased ratio of 12α-OH/non-12α-OH BAs. In male mice, this is associated with resistance to high fat diet-induced obesity and reduced intestinal lipid absorption. Therefore, data from this study (i) does not support the purported anti-obese effect of MCA and (ii) suggest that the 12α-OH BA content of the BA pool overrules hydrophobicity in the regulation of lipid absorption.

## AUTHOR CONTRIBUTIONS

Antwi-Boasiako Oteng designed the study, performed experiments, analyzed the data, and drafted the manuscript. Sei Higuchi performed experiments. Alexander S. Banks supervised experiments and analyzed data. Rebecca A. Haeusler designed the study, provided supervision, and revised the manuscript. All authors have read, edited, and approve of the final manuscript.

## ACKNOWLEDGMENTS/GRANT SUPPORT

This work was supported by NIH grants R01 DK115825 to RAH, T32 DK07559 to A-BO, S10OD028635 to ASB, and facilities/instrumentation supported by UL1TR001873, P30DK063608, P30DK026687, and American Diabetes Association grant 7-20-IBS-130. We are grateful to Utpal Pajvani and other members of the Haeusler Lab for discussions of the data.

## Abbreviations

12α-OH BA: 12-alpha hydroxylated bile acid
ALT: alanine transaminase
AST: aspartate transaminase
BA: bile acid
CM: chylomicron
Cyp2c: Cytochrome P450 2C
Cyp7a1: cholesterol 7-alpha-hydroxylase
Cyp7b1: 25-hydroxycholesterol 7-alpha-hydroxylase
Cyp8b1: sterol 12-alpha-hydroxylase
Cyp27a1: Cytochrome P450, family 27, subfamily a, polypeptide 1
CA: cholic acid
CDCA: chenodeoxycholic acid
DCA: deoxycholic acid
FXR: farnesoid X receptor
HDL: high density lipoprotein
HDCA: hyodeoxycholic acid
LCA: lithocholic acid
LDL: low density lipoprotein
MCA: muricholic acid
SHP: small heterodimer partner
UDCA: ursodeoxycholic acid
VLDL: very low density lipoprotein

## Graphical Summary

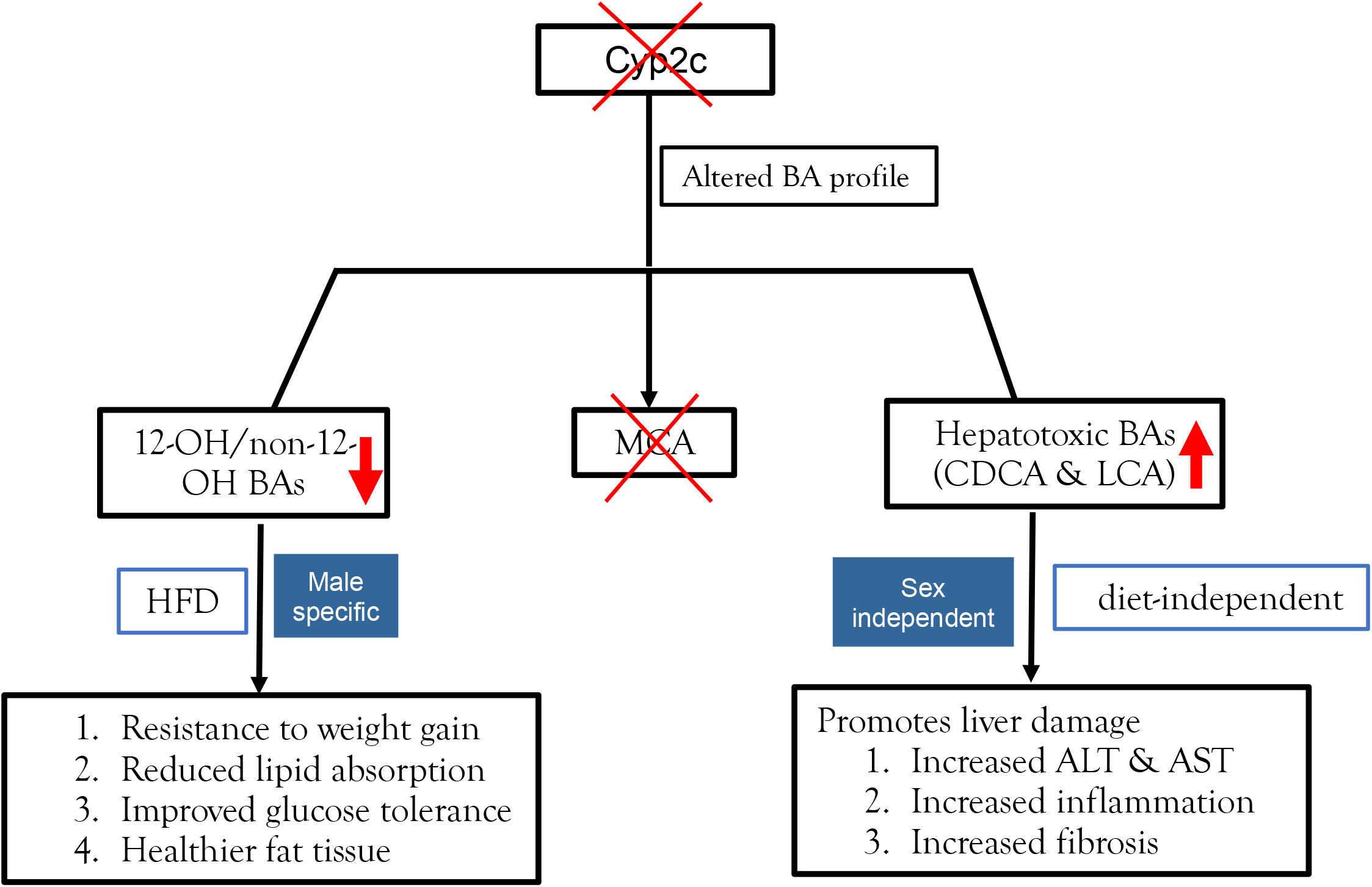

