## Supplementary data for "Absence of muricholic acid due to Cyp2c-deficiency protects against high fat diet-induced obesity in male mice but promotes liver damage"

Supplemental Table 1: Primer sequences of genes

| Genes | Forward primer | Reverse primer |
|---|---|---|
| 36b4 | ATGGGTACAAGCGCGTCCTG | GCCTTGACCTTTTCAGTAAG |
| Cyp2c-WT | CTACAATGCTCTGCCTACCC | AAATCTGACTCCCTCTTCTGG |
| Cyp2c-KO | GACATTGACATCCACTTTGCC | GATGGATGTGTGGAATGAAGAG |
| Cyp2c70 | TCCCAAGGGCACAAGTGTAAT | GCTGTAAGATGTTGGTCAGGATT |
| Cyp2c29 | ATCTGGTCGTGTTCCTAGCG | AGTAGGCTTTGAGCCCAAATAC |
| Cyp2c50 | ACTGTGGTGTTGCATGGATATG | GAGAAGCGCCTTGTGTTTTTC |
| Cyp2c67 | TGTGGGACTTATCCAGTGGC | ACCAAGTACAGTGCATGGGC |
| Cyp2c69 | TGTAGTCTTGGTGCTTTGTCTG | ACAATAGGCTGTGAGCCAAAATA |
| Cyp2c68 | GGGACTTATCCAGTGGCTCC | CAGCCAAGGTTCCAGCAATG |
| Cyp2c37 | AAACCAATGGCTCACCCTGT | GGGCTGCTCAGAATCTTTGT |
| Cyp2c54 | AGACAGAGCTATGAAAGAGGGAA | GTGAGAAGTGCCTCGTGTTTT |
| Cyp2c40 | GGCTCACAGCCTATTGTGGTA | TCAAAAACCGGAATCCTTCCTC |
| Cyp2c39 | GAGGAAGCATTCCAATGGTAGAA | TGTGAAGCGCCTAATCTCTTTC |
| Cyp2c38 | CACGGCCCATTGTTGTATTGC | TGAACCGTCTTGTCTCTTTCCA |
| Cyp2c55 | AATGATCTGGGGGTGATTTTCAG | GCGATCCTCGATGCTCCTC |
| Cyp2c65 | CATGGGGAGGAGTTTGCTGG | TGAAAACAACCCCGCAGTTT |
| Cyp2c66 | AGCCTGCTGTGGTGTTACAT | GCTGAAAACAACCCCACATCTT |
| Cyp2c44 | GCTGCCCTATACAGATGCCG | GTGACGCTAAGAGTTGCCCA |
| F4/80 | TGACTCACCTTGTGGTCCTAA | CTTCCCAGAATCCAGTCTTTCC |
| Cd68 | TGTCTGATCTTGCTAGGACCG | GAGAGTAACGGCCTTTTTGTGA |
| Mcp1 | TTAAAAACCTGGATCGGAACCAA | GCATTAGCTTCAGATTTACGGGT |
| Tnfa | CAACCTCCTCTCTGCCGTCAA | TGACTCCAAAGTAGACCTGCCC |
| Il6 | CTTCCATCCAGTTGCCTTCTTG | AATTAAGCCTCCGACTTGTGAAG |
| Srebp1c | GGAGCCATGGATTGCACATT | CCTGTCTCACCCCCAGCATA |
| Fasn | GGAGGTGGTGATAGCCGGTAT | TGGGTAATCCATAGAGCCCAG |
| Acaca | ATGGGCGGAATGGTCTCTTTC | TGGGGACCTTGTCTTCATCAT |
| Scd1 | TTCTTGCGATACACTCTGGTGC | CGGGATTGAATGTTCTTGTCGT |
| Dgat1 | TCCGTCCAGGGTGGTAGTG | TGAACAAAGAATCTTGCAGACGA |
| Col1a1 | GCTCCTCTTAGGGGCCACT | CCACGTCTCACCATTGGGG |
| Col1a2 | GTAACTTCGTGCCTAGCAACA | CCTTTGTCAGAATACTGAGCAGC |
| Timp1 | GCAACTCGGACCTGGTCATAA | CGGCCCGTGATGAGAAACT |
| aSma | GTCCCAGACATCAGGGAGTAA | TCGGATACTTCAGCGTCAGGA |
| Mmp2 | CAAGTTCCCCGGCGATGTC | TTCTGGTCAAGGTCACCTGTC |
| Fxr | GCTTGATGTGCTACAAAAGCTG | CGTGGTGATGGTTGAATGTCC |
| Cyp7a1 | GGGATTGCTGTGGTAGTGAGC | GGTATGGAATCAACCCGTTGTC |
| Cyp8b1 | CCTCTGGACAAGGGTTTTGTG | GCACCGTGAAGACATCCCC |
| Cyp27a1 | GCCTCACCTATGGGATCTTCA | TCAAAGCCTGACGCAGATG |
| Cyp7b1 | GGAGCCACGACCCTAGATG | TGCCAAGATAAGGAAGCCAAC |
| Shp | CGATCCTCTTCAACCCAGATG | AGGGCTCCAAGACTTCACACA |
| Srebp2 | CTGCAGCCTCAAGTGCAAAG | CAGTGTGCCATTGGCTGTCT |
| Hmgcr | AGCTTGCCCGAATTGTATGTG | TCTGTTGTGAACCATGTGACTTC |
| Ldlr | GCATCAGCTTGGACAAGGTGT | GGGAACAGCCACCATTGTTG |
| Pcsk9 | GAGACCCAGAGGCTACAGATT | AATGTACTCCACATGGGGCAA |
| Ucp1 | CAGCTTTGCCTCACTCAGGA | AAGCATTGTAGGTCCCCGTG |
| Dio2 | CTTCCTGGCGCTCTATGACTC | CCCCATCAGCGGTCTTCTC |
| Cox4i1 | CCGTCTTGGTCTTCCGGTTG | ACACTCCCATGTGCTCGAAG |

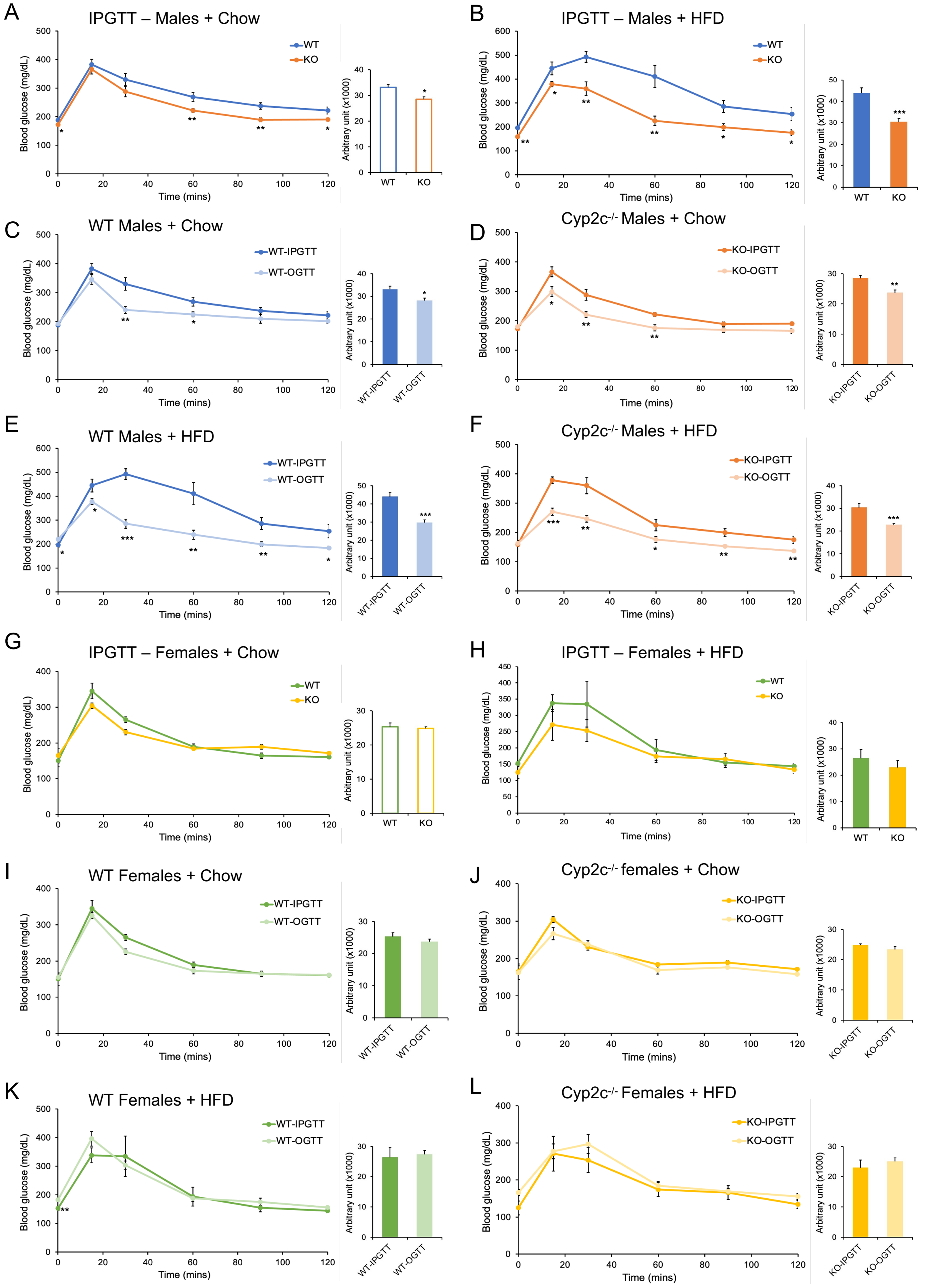


**Supplemental Figure 1.** Changes in glucose homeostasis is independent of incretin effect. A-B: Blood glucose level during IPGTT in chow-fed (A) and HFD-fed (B) male mice C-D: Comparison between IPGTT and OGTT for incretin effect in male WT (C) and male Cyp2c$^{-/-}$ (D) mice on chow diet. E-F: Comparison between IPGTT and OGTT for incretin effect in male WT (E) and male Cyp2c$^{-/-}$ (F) mice on HFD. G-H: Blood glucose level during IPGTT in chow-fed (G) and HFD-fed (H) female mice. I-J: Comparison between IPGTT and OGTT for incretin effect in female WT (I) and female Cyp2c$^{-/-}$ (J) mice on chow diet. K-L: Comparison between IPGTT and OGTT for incretin effect in female WT (K) and female Cyp2c$^{-/-}$ (L) mice on HFD. Insets represent area under the curve. Data are represented as mean ± SEM. n=7 mice/group for all groups, except n=5 mice/group for females + HFD. *p< 0.05, **p< 0.01 and ***p<0.001 relative to WT by Student's t-test.

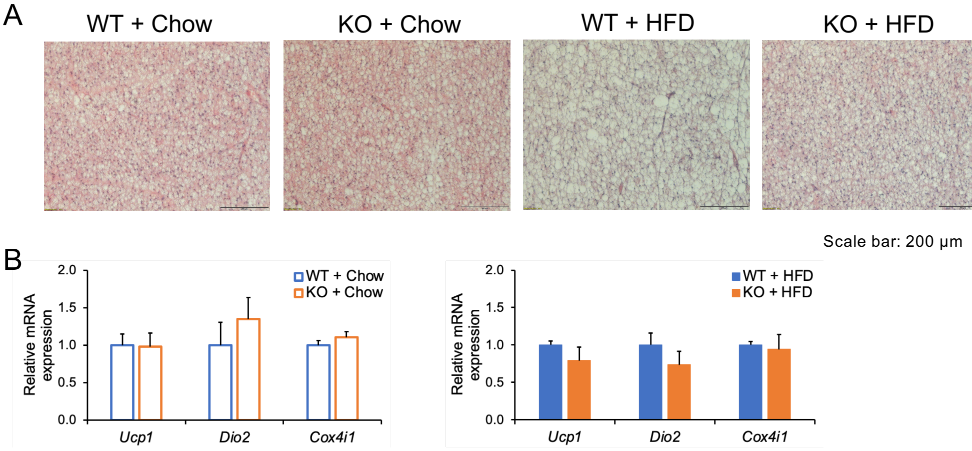


**Supplemental Figure 2.** No changes in BAT histology and gene markers of thermogenesis in male mice. A: Representative images of H & E staining of BAT. Scale bar is 200µm. B-C: mRNA expression of thermogenic genes in BAT of WT and Cyp2c$^{-/-}$ mice fed chow (B) or HFD (C) diets. Data are represented as mean ± SEM. n=7 mice/group. *$p < 0.05$, **$p < 0.01$ and ***$p < 0.001$ relative to WT by Student's t-test.
